# Computational redesign of the *Escherichia coli* ribose-binding protein ligand binding pocket for 1,3-cyclohexanediol and cyclohexanol

**DOI:** 10.1101/559328

**Authors:** Diogo Tavares, Artur Reimer, Shantanu Roy, Aurélie Joublin, Vladimir Sentchilo, Jan Roelof van der Meer

## Abstract

Bacterial periplasmic-binding proteins have been acclaimed as general biosensing platform, but their range of natural ligands is too limited for optimal development of chemical compound detection. Computational redesign of the ligand-binding pocket of periplasmic-binding proteins may yield variants with new properties, but, despite earlier claims, genuine changes of specificity to non-natural ligands have so far not been achieved. In order to better understand the reasons of such limited success, we revisited here the *Escherichia coli* RbsB ribose-binding protein, aiming to achieve perceptible transition from ribose to structurally related chemical ligands 1,3-cyclohexanediol and cyclohexanol. Combinations of mutations were computationally predicted for nine residues in the RbsB binding pocket, then synthesized and tested in an *E. coli* reporter chassis. Two million variants were screened in a microcolony-in-bead fluorescence-assisted sorting procedure, which yielded six mutants no longer responsive to ribose but with 1.2-1.5 times induction in presence of 1 mM 1,3-cyclohexanediol, one of which responded to cyclohexanol as well. Isothermal microcalorimetry confirmed 1,3-cyclohexanediol binding, although only two mutant proteins were sufficiently stable upon purification. Circular dichroism spectroscopy indicated discernable structural differences between these two mutant proteins and wild-type RbsB. This and further quantification of periplasmic-space abundance suggested most mutants to be prone to misfolding and/or with defects in translocation compared to wild-type. Our results thus affirm that computational design and library screening can yield RbsB mutants with recognition of non-natural but structurally similar ligands. The inherent arisal of protein instability or misfolding concomitant with designed altered ligand-binding pockets should be overcome by new experimental strategies or by improved future protein design algorithms.

---

Periplasmic binding proteins (PBPs) form a versatile superfamily of proteins with a conserved protein structure, named the bilobal structural fold^1,2^. PBPs facilitate nutrient and trace mineral scavenging for bacterial cells, by binding the ligand in the periplasmic space at high affinity and delivering the bound-ligand to a specific membrane-spanning transport channel^2^. Some PBPs are additionally involved in chemotactic sensing and interact in the ligand-bound state with a membrane-located chemoreceptor^3^. The crystal structures of several PBPs have been determined, showing two domains connected by a hinge region, with the binding pocket located between the two domains^3^. Both structure and nuclear-magnetic resonance data indicate that PBPs switch between two semi-stable conformations. Without ligand the protein adopts an *open* conformation, in which the binding site is exposed. Suitable ligand molecules become buried within the surrounding protein, and stabilize the *closed* protein form^4,5^. High-quality crystal structures of various PBPs have been determined, and this triggered pioneering ideas more than a decade ago to deploy PBPs as a generalized platform for computational design-based construction of new ligand-binding properties^6^. PBPs form an interesting class of proteins for biosensing. Biosensing can be achieved by measuring the intermolecular motion of the purified protein itself upon interaction with the target ligand^7^. Alternatively, the PBP protein is expressed in a living bacterial cell and triggers a synthetic signaling cascade upon ligand binding. This principle is embedded in so-called bioreporter cells or bactosensors^8^. By maintaining a single unique signaling cascade and reporter output, but varying the PBP-element with different ligand recognition, one could potentially develop a wide class of applicable bioreporters.

The concept of computational design of PBP variants with novel ligand-binding properties was proposed over a decade ago by the group of Hellinga and coworkers^9^. On the examples of five different PBPs in *Escherichia coli* they predicted and constructed mutant variants with binding pockets accommodating the non-natural substrates trinitrotoluene (TNT), lactate or serotonin at reported nM-mM *in vitro* affinities^9^. Particularly mutants of the ribose binding protein (RbsB) for TNT were further embedded in an *E. coli* synthetic bioreporter, in which ligand-bound RbsB-mutant contacts the Trz1 hybrid membrane receptor, increasing expression of a reporter gene fused to the *ompC* promoter^9^. This Trz1 receptor consists of a fusion of the 230 C-terminal amino acids of the *E. coli* EnvZ osmoregulation histidine kinase to the 265 N-terminal amino acids of the Trg methyl-accepting chemotaxis receptor protein^10^. Contact activation of Trz1 by ligand-bound RbsB triggers autophosphorylation of the cytoplasmic EnvZ-domain, leading to subsequent phosphorylation of the cognate response regulator OmpR, which activates the *ompC* promoter^11^. Independent engineering of the most sensitive published RbsB mutant (named TNT.R3), however, failed to reproduce the reported TNT detection at sub-μM concentrations in the *E. coli* Trz1-OmpR background and also failed to demonstrate TNT binding by a purifed TNT.R3 mutant using *in vitro* microcalorimetry^12^. Subsequent analysis of effects of alanine-substitutions in wild-type *E. coli* RbsB showed that mutations at 12 positions result in misfolded or poorly translocated proteins, one of which was also targeted in the TNT.R3 variant^13^. Purification and biophysical analysis of a further set of published mutant PBPs also failed to reproduce the original measurements, and suggested the cause being their misfolding and unintended oligomerization^14^. The initial studies may thus have underestimated to a large extent the propensity of PBPs to become misfolded as a result of binding pocket mutations.

More recently, ligand-specificities have been successfully interchanged between PBPs by using binding-pocket grafting (i.e., exchange of binding pockets between functionally closely related PBPs)^15,16^, improved prediction of native ligand binding^17^ and statistical coupling analysis (i.e., the prediction of mutations based on correlating amino acid residues in sectors of two classes of related proteins)^15^. So far, however, there have been no reports of non-cognate altered ligand-binding properties of PBPs. The goal of the underlying work was thus to revisit the concept of computational prediction of altered ligand-binding in RbsB. Because of the apparent difficulties to predict structure-function related side-effects such as protein folding, we hypothesized that predictions of minor changes in ligand-specificity might be more successful than major ones (e.g., from ribose to TNT). We thus chose to target molecules structurally related to ribose, in particular, 1,3-cyclohexanediol (13CHD) and cyclohexanol (CH). The computational protein design was based on exploration of sequence-space and estimations of Free energy of binding using Rosetta^18–20^, to identify a list of mutated protein sequences with potentially sufficiently low energy of binding with the new target ligands. The DNA encoding for a large set of approximately 2 million mutants was then chemically synthesized and cloned into a vector for screening of inducible GFP expression in the *E. coli* Trz1-OmpR, *ompCp::gfp* signaling reporter background^12^. Mutant libraries were screened on bead-encapsulated microcolony-grown cells by flow cytometry and sorted using fluorescence-assisted bead-sorting (FABS) (Fig. 1). Positively-responding mutant strains were recovered, their RbsB mutant proteins were purified and further characterized for *in vitro* ligand binding by isothermal microcalorimetry. Periplasmic abundance of the mutant proteins was quantified by peptide mass-spectrometry in comparison to wild-type RbsB, and their folding was addressed by circular dichroism spectroscopy. We recovered a small number of mutants with modest inducibility but significant change in ligand-binding specificity compared to wild-type RbsB and ribose, indicating that the computational design correctly targeted the intended new ligand-binding properties. However, despite some of gain of inducibility, our results indicate the mutant proteins to be unstable, and prone to misfolding during synthesis and secretion.

**Figure 1.**
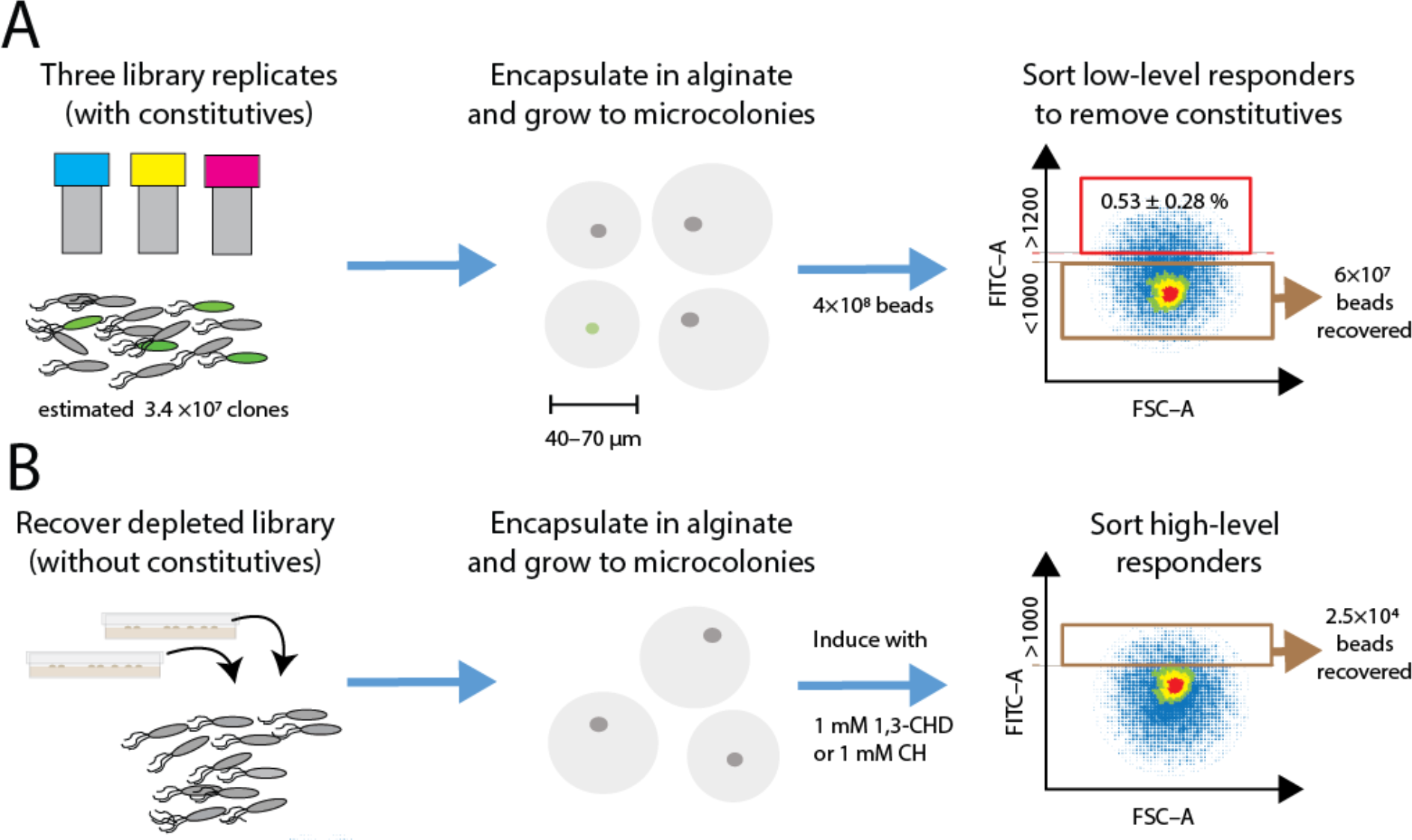
Overview of the mutant library screening strategy. A) Library replicates are encapsulated as on average one cell per agarose bead and grown to microcolony size in fumarate medium without inducer. Beads are passed on flow cytometry and beads with GFP fluorescence intensity below 1000 U are sorted and recovered. B) Cells are recovered from collected beads, again encapsulated and grown to microcolonies, after which they are exposed to the new ligands. Beads with GFP fluorescence intensity higher than 1000 U are recovered and further screened.

## Results

### RbsB mutant library design

Exploration of sequence space using Rosetta enzyme design simulations produced a library of targeted RbsB mutants, which were predicted to have improved affinity for the non-cognate ligands 13CHD and/or CH. The used design template was the scaffold of wild-type ribose-binding protein RbsB of *E. coli* in its closed configuration (PBD ID: 2DRI, Fig. 2A). First, prior to the design simulations, interactions between RbsB and ribose, 13CHD or CH were studied using docking and molecular dynamics simulations in CHARMM and Merck Molecular Force Fields (MMFF), to gain intuition on the stability of the binding pocket. The average spatial deviation of the RbsB binding pocket with placed ribose during 2 ns (as the root-mean squared deviation) was less than 0.2 Å, but for docked 13CHD and CH molecules was around 0.3 Å. The ligands themselves showed varying positions with an average root-mean squared deviation of 0.7 Å for ribose, 1 Å for 13CHD and 1.5 Å for CH. Calculated free energies of binding (ΔG_binding_) from CHARMM and MMFF were lowest for ribose, as expected, with −38.35 kcal mol^−1^, but −19.57 kcal mol^−1^ for 13CHD and −14.31 kcal mol^−1^ for CH. These results indicated unstable interactions of 13CHD and CH in the wild-type RbsB binding pocket. We conducted a per-residue free energy decomposition analysis21 using the simulation trajectories. The poorer ΔG_binding_ of 13CHD and CH seemed largely contributed by the RbsB residues D89, R90 and D215 (for 13CHD), and D89, D191 and D215 (for CH) (Fig. S1).

**Figure 2.**
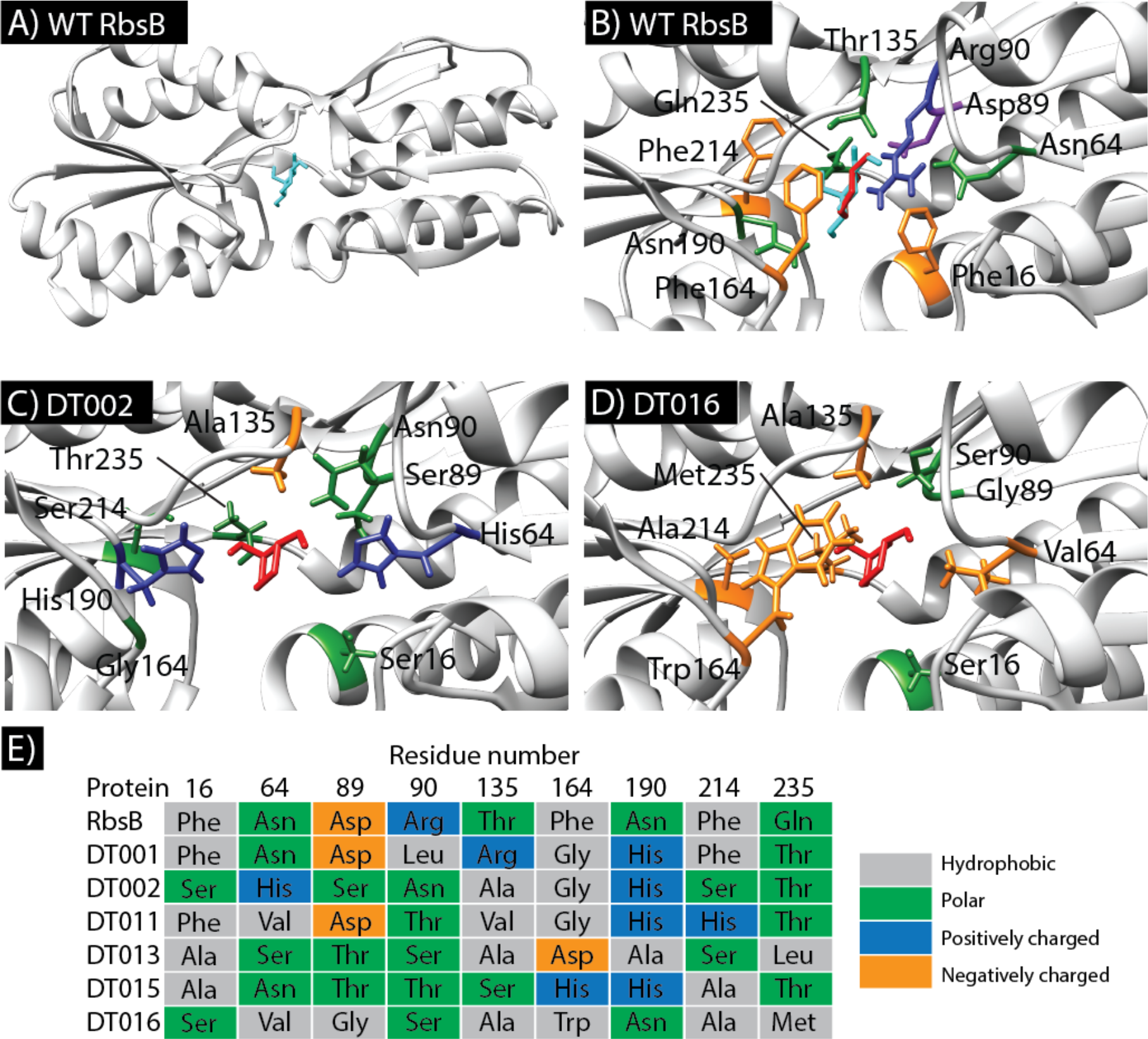
Structures of wild-type RbsB (WT RbsB) and DT002, DT016 mutants. (A) Global view of closed RbsB (PDB ID: 2DRI) molecular structure with ribose (cyan) bound in its pocket. (B) Details of the RbsB binding pocket with 13CHD (red) and ribose (cyan) molecules. Critical amino acid residues for substrate binding are indicated and color-coded based on amino acid characteristics (nonpolar-orange; positively charged-blue; polar-green; negatively charged-purple). (C) Details of the DT002 binding pocket (threaded on the RbsB structure, PDB ID: 2DRI) with 13CHD (red, placed according to docking with Rosetta) with indication of mutated amino acids. (D) Same as in (C) but for the DT016 mutant. (E) Overview of the targeted residues in the recovered mutants compared to wild-type RbsB.

Next, we used Rosetta to predict potential beneficial mutations in RbsB for binding of 13CHD and CH. Defined key residues in the RbsB binding pocket (Table 1, Fig. 2B) were computationally replaced by alanine. The 13CHD and CH molecules were then docked 1000 times independently into the ‘’stripped” binding pocket to obtain the positions with the lowest minimal binding energy, which were used as a starting point for the design mode. Design simulations (100 repetitions each producing 100 designs) then explored the combinatorial mutations on the defined 9 positions, from which pool 200 sequences were ranked according to the lowest predicted binding energy, minimal packing energy, hydrogen bond counts, and ligand-solvent exposure. This yielded a list of one of four possible amino acids at the 9 positions in RbsB (Table 1). The DNA encoding these RbsB variants in all their combinations plus the original wild-type residue was produced as a mixed library by DNA synthesis, cloned and introduced into an *E. coli* host enabling GFPmut2 production through the hybrid Trz1-OmpR signaling chain (strain 4172, Table 2, Fig. S2). Independent cloning reactions resulted in three mutant libraries with estimated sizes of 7×10^6^, 24×10^6^ and 3.3×10^6^ primary transformants.

**Table 1.** Mutations introduced into *E. coli* wild-type RbsB protein

| Amino acid | Wild-type residue | Tested mutations |
| --- | --- | --- |
| 16 | F | H, T, S, A |
| 64 | N | V, I, H, A |
| 89 | D | G, T, S, V |
| 90 | R | S, N, T, L |
| 135 | T | S, A, R, V |
| 164 | F | D, G, H, W |
| 190 | N | H, S, T, A |
| 214 | F | H, A, S, N |
| 235 | Q | T, D, L, M |

**Table 2.** List of strains used in this study

| Strain | Host | Plasmid(s) | Relevant characteristics | Reference or source |
| --- | --- | --- | --- | --- |
| 97 | *E. coli* BL21 (DE3) | pLysS | Host strain for overexpression from the T7 promoter | <sup>35</sup> |
| 3725 | *E. coli* BL21 (DE3) | pLysS, pAR1 | Cytoplasmic overexpression of RbsB-His<sub>6</sub> | This work |
| 4172 | *E. coli* BW25113 ΔrbsB | pSYK1 | Host strain containing the P<sub>tac</sub>-trz1, P<sub>ompC</sub>-gfpmut2 bioreporter system | <sup>12</sup> |
| 4175 | *E. coli* BW25113 ΔrbsB | pAR3, pSYK1 | Expression of RbsB with signal sequence for periplasmic translocation | <sup>12</sup> |
| 5913 | *E. coli* BW25113 ΔrbsB | pSTV-DT001, pSYK1 | As 4175, but for DT001 mutant protein of RbsB | This work |
| 5903 | *E. coli* BW25113 ΔrbsB | pSTV-DT002, pSYK1 | As 4175, but for DT002 mutant protein of RbsB | This Work |
| 5904 | *E. coli* BW25113 ΔrbsB | pSTV-DT011, pSYK1 | As 4175, but for DT01 mutant protein of RbsB | This Work |
| 5905 | *E. coli* BW25113 ΔrbsB | pSTV-DT013, pSYK1 | As 4175, but for DT013 mutant protein of RbsB | This Work |
| 5906 | *E. coli* BW25113 ΔrbsB | pSTV-DT015, pSYK1 | As 4175, but for DT015 mutant protein of RbsB | This Work |
| 5907 | *E. coli* BW25113 ΔrbsB | pSTV-DT016, pSYK1 | As 4175, but for DT016 mutant protein of RbsB | This Work |
| 5999 | *E. coli* BW25113 ΔrbsB | pSTV-DT016 (W164G), pSYK1 | As 4175, but for DT016<sub>W164G</sub> mutant protein of RbsB | This Work |
| 6054 | *E. coli* BW25113 ΔrbsB | pSTV-DT016 (M235V), pSYK1 | As 4175, but for DT016<sub>M235V</sub> mutant protein of RbsB | This Work |
| 5927 | *E. coli* BL21 (DE3) | pET3d-DT001, pLysS | Cytoplasmic overexpression of DT001-His<sub>6</sub> | This Work |
| 5908 | *E. coli* BL21 (DE3) | pET3d-DT002, pLysS | Cytoplasmic overexpression of DT002-His<sub>6</sub> | This Work |
| 5909 | *E. coli* BL21 (DE3) | pET3d-DT011, pLysS | Cytoplasmic overexpression of DT011-His<sub>6</sub> | This Work |
| 5910 | *E. coli* BL21 (DE3) | pET3d-DT013, pLysS | Cytoplasmic overexpression of DT013-His<sub>6</sub> | This Work |
| 5911 | *E. coli* BL21 (DE3) | pET3d-DT015, pLysS | Cytoplasmic overexpression of DT015-His<sub>6</sub> | This Work |
| 5912 | *E. coli* BL21 (DE3) | pET3d-DT016, pLysS | Cytoplasmic overexpression of DT016-His<sub>6</sub> | This Work |
| 6016 | *E. coli* BL21 (DE3) | pET3d-DT016(W164G), pLysS | Cytoplasmic overexpression of DT016<sub>W164G</sub>-His<sub>6</sub> | This Work |

A total of 4×10^8^ alginate beads encapsulating individual cells from the mutant libraries and grown to microcolonies was screened by FABS for GFPmut2 fluorescence, in first instance in absence of inducer (Fig. 1A). An estimated 0.53 ± 0.28 % of the screened beads displayed fluorescence above 1200 units under non-induced conditions, and were considered constitutive-ON mutants. Approximately 60 million beads with fluorescence below 1000 units were sorted and recovered as mixture. Cells were released from the beads, freshly cultured, encapsulated in new alginate beads, regrown to microcolonies and induced with 1 mM 13CHD (Fig. 1B). In this second phase, beads with a fluorescence level higher than 1000 units were sorted and plated to grow individual colonies (a total of 2.3×10^4^). After rescreening six mutants displayed consistently between 1.2-1.5-fold higher GFP fluorescence upon incubation with 1 mM 13CHD in comparison to media alone, which was a moderate response but statistically significant (p-values < 0.05). These mutants were no longer inducible and even slightly inhibited with 0.1 mM ribose (Table 3). In contrast, the same *E. coli* host expressing wild-type RbsB was not inducible with 13CHD but is 13-fold inducible with 0.1 mM ribose (Table 3). Only one of the six mutants (DT016) responded to 1 mM CH with a statistically significant increase in GFP fluorescence (Table 3).

**Table 3.** GFPmut2 induction in *Escherichia coli* expressing either wild-type or mutant RbsB.

| Protein | GFPmut2 uninduced fluorescence | Fold induction<sup>a</sup> |  |  |
| --- | --- | --- | --- | --- |
|  |  | Ribose<sup>b</sup> | 1,3-Cyclohexanediol<sup>b</sup> | Cyclohexanol<sup>b</sup> |
| **Wild-type** | 6180 ± 1580<sup>c</sup> | **13 ± 2.2<sup>d</sup>** | **0.8 ± 0.03** | **0.8 ± 0.05** |
| **DT001** | 6846 ± 1658 | **0.9 ± 0.05** | **1.3 ± 0.16** | 0.9 ± 0.13 |
| **DT002** | 6019 ± 886 | 1.0 ± 0.05 | **1.4 ± 0.13** | 0.9 ± 0.08 |
| **DT011** | 8462 ± 440 | 0.9 ± 0.09 | **1.2 ± 0.04** | **0.8 ± 0.05** |
| **DT013** | 6154 ± 1319 | 0.9 ± 0.09 | **1.3 ± 0.6** | **0.8 ± 0.08** |
| **DT015** | 4481 ± 694 | **0.9 ± 0.16** | **1.3 ± 0.12** | 0.8 ± 0.11 |
| **DT016** | 24430 ± 3460 | 1.0 ± 0.08 | **1.5 ± 0.01** | **1.5 ± 0.05** |
| **DT002<sub>H190N</sub>** | 5054 ± 805 | 0.9 ± 0.07 | 0.9 ± 0.15 | **0.7 ± 0.03** |
| **DT016<sub>W164G</sub>** | 11627 ± 2028 | **0.8 ± 0.12** | **1.3 ± 0.04** | **1.3 ± 0.01** |
| **DT016<sub>M235V</sub>** | 5960 ± 1300 | **0.8 ± 0.09** | 1.2 ± 0.08 | 1.2 ± 0.06 |

a) Mean GFPmut2 fluorescence in the assay with inducer divided by that of the assay with buffer only. Assay incubation time is 2h30 at 37°C. Values from eight replicates, ± calculated SD.
b) Final concentration of inducers in the assay: ribose, 0.1 mM; 13CHD, 1 mM; CH, 1 mM.
c) Mean values from eight replicates, ± calculated SD.
d) Values in bold indicate statistically significant induction (pair-wise T-test; p <0.05, one-sided, equal variance).

All six recovered mutants contained different amino acid substitutions, with some, but little overlap (Fig. 2E). Mutant DT001 displayed five mutations and four wild-type residues at the 9 targeted positions, followed by DT011 with 7 mutations, DT015 and DT016 with 8, and clones DT002 and DT013 with all 9 targets substituted (Fig. 2E). In the majority of the 13CHD-responsive mutants, positions D89, R90 and Q235 were replaced by a polar residue, whereas position T135 was replaced by a non-polar residue. Also, in four of six mutants, N190 was substituted by a histidine (Fig. 2E).

### Reduced periplasmic space abundance of RbsB mutants responsive to 13CHD

The relative periplasmic space abundance of four RbsB mutants determined by quantitative mass spectrometry was lower compared to wild-type RbsB (Table 4). The DT002 and DT015 proteins displayed the lowest relative abundance, followed by DT011 and DT001. The periplasmic abundance of mutant DT016 was 2 times higher than RbsB. Interestingly, the relative abundance of MglB (galactose-binding protein) was higher in the periplasmic space of *E. coli* expressing mutants DT011 or DT015, in comparison to those expressing wild-type RbsB or the other mutant proteins (Table 4). Also, the summed abundance of all periplasmic binding proteins (excluding RbsB) was higher in all *E. coli* expressing RbsB mutant proteins than wild-type, with up to between 4.5 and 6 times increase in mutants DT015 and DT011 (Table 4). Quantitative mass spectrometry data thus suggest that translocation was affected for most RbsB mutant proteins and that this also influenced the translocation of other periplasmic binding proteins to the periplasm.

**Table 4.** Periplasmic abundance of wild-type or mutant RbsB proteins in *Escherichia coli*.

| Expressed protein | Periplasmic abundance |  |  |  |  |  |
| --- | --- | --- | --- | --- | --- | --- |
|  | Exclusive peptide count<sup>a</sup> |  |  | Normalized count<sup>b</sup> |  |  |
|  | (mutant) RbsB | MglB<sup>c</sup> | Periplasmic binding proteins<sup>d</sup> | (mutant) RbsB | MglB<sup>c</sup> | Periplasmic binding proteins<sup>d</sup> |
| RbsB | 115 | 46 | 91 | 72 | 29 | 57 |
| DT001 | 56 | 38 | 235 | 46 | 31 | 193 |
| DT002 | 13 | 7 | 55 | 24 | 13 | 101 |
| DT002<sub>H190N</sub> | 59 | 38 | 301 | 53 | 34 | 268 |
| DT011 | 39 | 161 | 348 | 42 | 175 | 379 |
| DT015 | 24 | 202 | 407 | 16 | 133 | 268 |
| DT016 | 143 | 19 | 141 | 181 | 24 | 179 |
| DT016<sub>W164G</sub> | 148 | 20 | 183 | 159 | 21 | 197 |

### *In vitro* 13CHD binding by mutant RbsB

Cytoplasmic overexpressed His_6_-tagged RbsB was readily purified and resulted in protein with >97% purity on SDS-PAGE and a molecular mass of around 30 kDa as expected (Fig. 3A, black triangle). Contrary to wild-type RbsB, contaminating proteins were consistently observed in purified His_6_-tagged mutant RbsBs. One or two prominent contaminants with a mass of around 70 kDa were observed after affinity (Fig. 3B and C, red triangle) and gel filtration columns. These contaminants contributed to an estimated 5-15 % of the total protein quantity. Interestingly, addition of 10 mM ATP to the eluted protein fraction after affinity purification but before gel filtration led to removal of these contaminants (Fig. 3D). Possibly, therefore, they consisted of *E. coli* chaperones such as Hsp70 or DnaK, involved in protein folding and refolding22, which remained attached to the mutant RbsBs and detached upon addition of ATP. This suggests that the mutant RbsB proteins produced in the *E. coli* cytoplasm suffer from partial misfolding and are stabilized by chaperones^23^.

**Figure 3.**
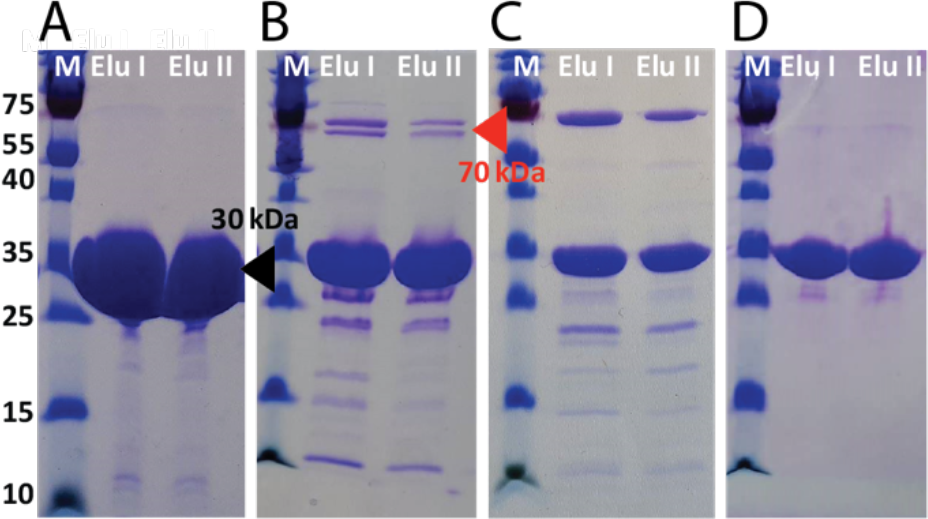
Overexpression and purification of RbsB-His_6_ and mutants DT002-His_6_ and DT016-His_6_. (A) SDS-PAGE gel of purification steps with a HisTrap column of RbsB-His_6_. (B) and (C) as for panel (A) but for mutants DT0016-His_6_ and DT002-His_6_, respectively. (D) SDS-PAGE gel of elution steps after a gel filtration column for mutant DT016-His_6_. M, Marker; Elu, Elution step. Black triangle indicates the expected position of RbsB-His_6_, DT002-His_6_ and DT016-His_6_ proteins. Red triangle indicates the position of the assumed *E. coli* chaperones.

Binding of ribose to wild-type purified RbsB-His_6_ in isothermal titration calorimetry (ITC) resulted in a clear heat release with an estimated binding affinity constant K_D_ of 530 nM (Fig. 4A), which is similar to literature values^14^. Purified RbsB-His_6_ showed no significant interaction with 13CHD (Fig. 4D). In contrast, modest but consistent heat release was observed with purified DT002-His_6_ and DT016-His_6_ in presence of 13CHD in comparison to buffer alone (Fig. 4B, C). Assuming binding of a single 13CHD ligand per protomer, we found an apparent K_D_ of 190 μM for DT002-His_6_ and 5 μM for DT016-His_6_. Kinetic heat release and molar ratios suggested that actually only part of the purified protein fraction engages in binding of the ligand, possibly because another fraction was misfolded and inactive (Fig. 4B, C). No binding of ribose by either of the two mutant proteins was observed (Fig. 4E, F), and none of the other purified mutant proteins (DT001-His_6_, DT011-His_6_, DT013-His_6_ or DT015-His_6_) yielded measurable heat release with either 13CHD or ribose as substrates (not shown).

**Figure 4.**
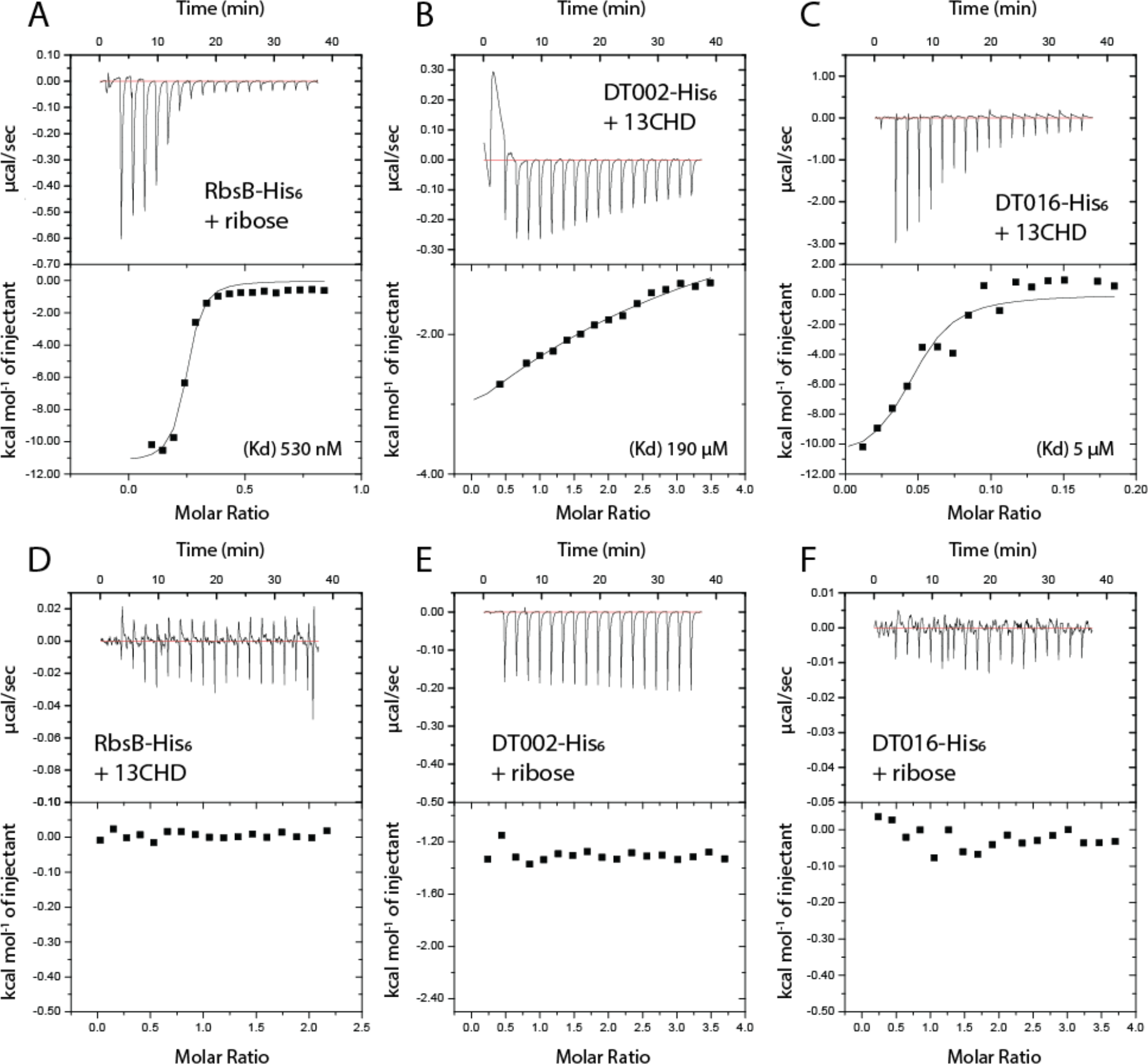
*In vitro* ligand binding measurements using isothermal microcalorimetry (ITC) with purified proteins. (A) Binding affinity of RbsB-His_6_ protein with ribose. (B) Binding affinity of DT002-His_6_ protein with 13CHD. (C) DT016-His_6_ protein with 13CHD. (D) RbsB-His_6_ protein with 13CHD. (E) DT002-His_6_ protein with ribose. (F) DT016-His_6_ protein with ribose. Kd, constant of affinity, assuming a single-ligand per protomer binding model. Graphs display immediate heat release in μcal s^−1^ (upper panels) and calculated heat released per mol of injectant (lower panels).

The RbsB protein fraction after affinity purification and gel filtration was stable and displayed consistent binding to ribose in ITC, even upon −80°C freezing and thawing of aliquoted fractions. In contrast, mutant protein fractions purified in the same manner except for addition of ATP before gel filtration were unstable. Purified DT002-His_6_ fractions could be kept on ice for at least 4 h and produced similar heat release in ITC upon addition of 13CHD for three consecutive measurements. In contrast, after freezing at −80°C and thawing, the apparent binding affinity was reduced and sometimes even lost. 13CHD-binding to purified DT016-His_6_ disappeared within 2 or 3 h after purification, even while maintaining the protein solution on ice. After −80°C freezing and thawing, the DT016-His_6_ protein fraction no longer showed any heat-release from added 13CHD in ITC. These observations and the poor molar ratio of 13CHD binding (Fig. 4B, C) suggested that the DT002-His_6_ and DT016-His_6_ mutant proteins have strongly reduced stability and spontaneously misfold during purification and ITC. Not unlikely, the other four RbsB mutant proteins already completely misfolded during purification, and no sufficiently stable fractions were obtained to measure productive ligand-binding in ITC.

### Secondary structure changes in mutant proteins compared to RbsB

To detect secondary structure differences between wild-type and mutant proteins and observe ligand-induced changes, we analyzed purified protein fractions by circular dichroism spectroscopy in absence and presence of ligand (ribose or 13CHD, Fig. 5). All three His-tagged proteins (RbsB, DT002 and DT016) had similar circular dichroism spectra but with different Δε intensities, which slightly (RbsB and DT002) or more importantly (DT016) increased upon addition of their ligands (Fig. 5A). Secondary structure protein-fold predictions from circular dichroism spectra using recently published tools^24^ on repeated independently purified protein batches indicated DT002 and DT016 to carry smaller proportions of helices but increased proportions of anti-parallel/parallel and ‘turn’-folds compared to RbsB (Fig. 5B). This suggests notable distortions in the RbsB-folds as a result of the introduced mutations (Fig. 2). Addition of ribose to RbsB resulted in a notable predicted reduction of the antiparallel-2 relaxed fold (for definition, see Ref^24^) and an increase of ‘other’ folds and turns (Fig. 5B). This might correspond to the closed configuration of the protein (see, e.g., Fig. 3B in Reimer *et al*.^13^). This decrease of the proportion of antiparallel-2 relaxed fold was also observed in one preparation of the purified DT016-protein after addition of 13CHD (see asterisk within Fig. 5B), but not with addition of ribose or CH. Addition of ligands to DT002 protein preparations did not cause any consistent or pronounced changes in the predicted secondary structure fold composition (Fig. 5B).

**Figure 5.**
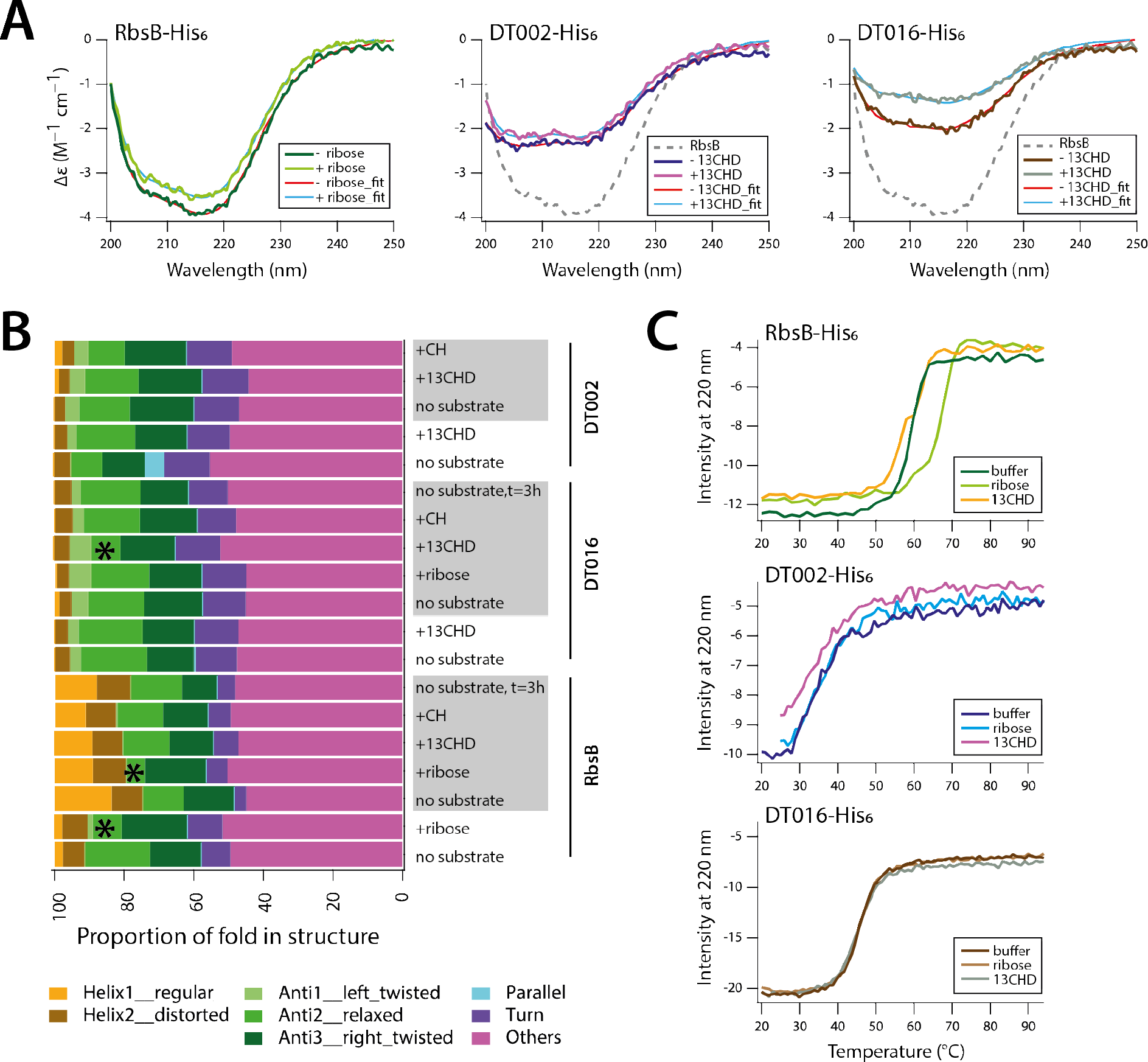
Secondary structure analysis and temperature melting curves of purified wild-type RbsB-His_6_, DT002-His_6_ and DT016-His_6_. (A) Circular dichroism spectra of purified proteins in buffer A (without imidazole) at a protein concentration of 0.1 mg ml^−1^, in absence or presence of inducer (ribose, 0.1 mM, CH or 13CHD, 1 mM). Spectra were fitted according to reference^24^. (B) Inferred secondary structure fold composition of the three purified proteins in two independent purified batches (alternating white or grey background), in presence or absence of ligands. Asterisks note the significant changes in antiparallel-2 relaxed protein fold upon productive ligand-addition. Protein fold terminology as in reference^24^. (C) Melting curves of purified proteins at a protein concentration of 0.3 mg ml^−1^, in buffer or in presence of ribose (0.1 mM) or 13CHD (1 mM).

Melting curves of wild-type, DT002 and DT016 purified protein fractions indicated important differences in thermal stability (Fig. 5C). Whereas wild-type RbsB showed a melting temperature (Tm) of 58.9±0.1°C (Sigmoidal curve fitting), that of DT016 was only 46.1±0.1°C and that of DT002 not more than 34.7±0.7°C (Fig. 5C). A robust shift in Tm of ~8°C was observed for RbsB upon addition of 0.1 mM ribose (Fig. 5C). This result is in accordance with previously reported data^14,25^. In presence of 1 mM 13CHD, however, the Tm of RbsB was slightly reduced to 58.5±0.2°C (Fig. 5C). In contrast, the melting temperatures of mutants DT016 and DT002 were not measurably affected by addition of ribose or 13CHD (Fig. 5C). This indicated that both DT002 and DT016 mutant proteins were indeed much less stable than wild-type RbsB and that interaction with 13CHD did not further stabilize the proteins.

### Effect of secondary mutations

In order to potentially improve the stability and/or translocation to the periplasm of the two most promising mutants (DT002 and DT016), further targeted mutations were introduced on these proteins. By site-directed mutagenesis H190 was reverted to wild-type N190 in mutant DT002 (Fig. 2E). Previous studies indicated the importance of N190 on RbsB stability and/or translocation^13,25^. In the mutant DT016 the residue 235, which had also been suggested to be implicated in RbsB stability^25^, was changed from M235 to V235. Position W164 in the DT016 protein, which is very close to the binding pocket and might block ligand access, was randomized (Fig. 2D). Back mutation of H190N in DT002 led to complete loss of inducibility by 13CHD (Table 3). Also the M235V mutation in mutant DT002 led to complete loss of 13CHD inducibility. Of the randomized positions at W164 in mutant DT016, only a replacement to Gly maintained 13CHD and CH induction (Table 3). The periplasmic abundance of DT002_H190N_ improved compared to DT002, whereas that of DT016_W164G_ remained the same as that of DT016 (Table 4).

## Discussion

PBPs have attracted wide interest because of their potential for biosensing and as universal scaffold for engineering ligand-binding properties *à la carte*^6^. However, despite detailed structure information on a number of PBPs^2^, and their biochemical, biophysical and genetic characterization, this *à la carte* design has remained largely elusive^14,26^. Structure-guided computational predictions to change ligand-binding specificities have been to some extent successful for other sensory-type proteins, such as transcription factors^26–28^, but for PBPs have remained limited to small modifications of binding properties in existing ligands^29^. Using the well-characterized RbsB protein from *E. coli* as a model, we showed here that computational predictions of altered amino acid residues in the RbsB binding pocket can indeed lead to a change of functional binding of the cognate substrate (ribose) to foreign but chemically related ligands (13CHD and CH). We acknowledge that although the loss of ribose-binding by the derived mutant RbsB proteins is very clear, the gain of new functionality is detectable but small. The very modest functional gain is not surprising and has been more frequently observed in similar library-screening efforts for altered PBP ligand-binding pocket designs^15^. Our data suggest that the main reason for the limited functional gain is the apparent propensity of RbsB to become misfolded upon mutational redesign of the binding pocket. So far, we have not been able to improve these mutants further by secondary mutations.

Several lines of evidence support our conclusion that we obtained a true change of cognate ligand-binding specificity of RbsB to a non-natural ligand, starting from Rosetta simulations and predictions of improved 13CHD- and CH-binding by changes in 9 amino acids. First of all, six different mutant proteins were isolated from the synthesized clone library, with up to 1.5-fold times induction with 13CHD in the *E. coli trz1-ompCp-gfpmut2* reporter strain. These mutants had lost completely the capacity to become induced *in vivo* by ribose, and RbsB as well as the majority of other mutants in the library showed no induction with 13CHD (Table 3). Further mutation of a number of altered residues in the mutants DT002 and DT016 resulted in loss of 13CHD induction (Table 3). Secondly, ITC measurements confirmed binding of 13CHD by the mutants DT002 and DT016, with estimated K_D_ of 190 μM and 5 μM, respectively (Fig. 4B, C). This is indicative for poor binding, but a K_D_ of 5 μM is in the range of measured affinity (1.6 μM) of a grafted L-glutamine-binding domain on the *Salmonella typhimurium* LAO periplasmic binding protein^15^. Finally, circular dichroism spectroscopy and secondary structure fold-decomposition using a recent new approach^24^ indicated structural changes to occur in DT016 upon 13CHD addition (Fig. 5B), although this was not consistently observed in independent protein preparations.

Our results corroborated previous observations that computationally designed PBP variants suffer from misfolding and instability^14,15^. Despite showing some inducibility in the *E. coli* signaling reporter chain, four out of six proteins failed to show 13CHD binding in ITC, likely because they unfolded during purification. The two most stable mutant proteins DT002 and DT016, quickly lost functional activity in ITC upon purification and were dramatically less stable than wild-type in thermal denaturation (Fig. 5C). All mutant proteins showed evidence for chaperone co-purification after affinity chromatography, indicative for misfolding. This obviously hampers the screening of mutant libraries, given that the procedure relies on functional gain-of-GFP fluorescence (Fig. 1). If we assume that the observed induction is a combination of the ‘true’ affinity of the mutant protein for its new ligand and the ratio of correctly versus misfolded protein, the actual gain of 13CHD- and CH-binding may be higher than an induction factor of 1.5 suggests.

Detecting and separating mutants from the initial library with such small improvement of inducibility by 13CHD required optimization of the screening method. We initially screened the library for gain-of-fluorescence upon induction on individual cells, but found that single cell variation was too high and resulted in many false-positive signals^30^. Instead, therefore, we switched to growing reporter cells inside alginate beads to microcolonies, which improved the reproducibility and screening efficiency, and reduced the number of false positives. Others have recently compared such procedures and have come to the same conclusion^31^. Surprisingly, around 0.5 % of the clones in our library were constitutively 'ON', showing high GFP fluorescence even in absence of inducer. Constitutives may be the result of combinations of mutations stabilizing the closed configuration without inducer being present. Once in the closed conformation the RbsB 'ON'-mutant may bind to the membrane receptor Trz1 and trigger the bioreporter system. The addition of a step to first sort mutants with lower fluorescence levels in absence of inducer was essential to remove the constitutive 'ON'-mutants and improve the screening efficiency (Fig. 1A).

What do the recovered mutants tell us about potential ligand-binding in their designed pockets? The ribose-binding pocket of RbsB has been investigated in detail in previous studies. Vercillo *et al.* reported 13 amino acid positions (S9, N13, F15, F16, N64, D89, S103, I132, F164, N190, F214, D215 and Q235) to play an important role in ribose binding^25^. Molecular dynamics simulations suggested several of those to be limiting 13CHD- or CH-binding by RbsB (Fig. S2). In the final computational strategy we stripped the presumed RbsB binding pocket at nine positions (changing virtually to Ala-residues), sampled the positions for 13CHD and CH with the lowest ΔG, and predicted the sets of amino acid residues to contribute with improved 13CHD and CH binding. Although this seemed the best strategy at the time, a more recent and complete Ala-substitution screening of RbsB by our group found four crucial residues for ribose recognition (D89, N190, D215, R141)^13^, two of which (D215, R141) were not included in the library predictions. In contrast, that study found several residues critical for RbsB folding and/or translocation, notably D89 and N190, which were targeted here. Indeed, in four out of the six isolated 13CHD-responsive mutants the D89 residue was substituted by a polar amino acid. The N190 residue was substituted in 5 out of the 6 mutants, of which four times by a histidine, a positively charged amino acid (Fig. 2E). This may thus have indirectly contributed to mutant proteins with poorer stability. Perhaps not surprisingly at this stage, several different new binding pocket configurations appeared to confer measurable gain-of-function of 13CHD binding and loss of ribose binding (Fig. 2E). These converge to some extent in the character of the newly positioned amino acid residue, but not in their exact type. Given that the modeled mutant protein binding pockets may in reality deviate more than is suggested in Figure 2C and D, it is too speculative to infer how the introduced new amino acid residues might be contributing to the binding of 13CHD.

Mass spectra analysis revealed that all 13CHD-responsive mutant proteins, except DT016 and its derivative DT016_W164G_, were less abundant in the periplasmic space than wild-type RbsB (Table 4). This is indirect evidence that the mutant proteins may have additional difficulties in translocation to the periplasm. Cells expressing DT011 and DT015 displayed higher periplasmic levels of MglB and other proteins (Table 4), which might be due to their increased flux through the Sec-translocation channel in absence of lesser abundant mutant RbsB. In case of cells expressing DT016 or DT016_W164G_, periplasmic space abundance of the mutant protein was higher but that of other PBPs was lower, perhaps because of competition with RbsB through the Sec-channel (Table 4). In case of DT002, both its own periplasmic space abundance, as well as that of MglB and other PBPs, were lower. This may be the result of a partial blocking of the translocation system by the DT002 protein. Four mutant proteins (DT001, DT002, DT011 and DT015) with lower abundance in the periplasm than wild-type RbsB carried an amino acid substitution at the N190 position. Ala-substitutions at this position resulted in loss of ribose-induction, potential misfolding and/or poor translocation into the periplasm^13,25^. Back mutation of the H190 residue in DT002 to an Asn, indeed increased its periplasmic space abundance (Table 4), but also resulted in loss of induction by 13CHD (Table 3). Mutant DT016, on the other hand, still retained asparagine at position 190 and its periplasmic space abundance was twice as high as wild-type RbsB protein (Table 4). Instead, we suspected that the bulky Trp-residue at position 164 in DT016 would limit protein flexibility in the entry and hinge regions (Fig. 2D), and perhaps be responsible for the high observed fluorescence background in absence of inducer (Table 3). Indeed, replacing the Trp by a Gly (DT016_W164G_) resulted in a much lower background, similar periplasmic space abundance (Table 4) and retainment of 13CHD and CH induction potential (Table 3).

Our results underscore that design of new ligand properties in highly flexible proteins such as PBPs is very challenging. Scoring functions developed for evaluation of protein-ligand binding free energies are not accurate enough^26,32^, and it is extremely difficult to predict the intrinsic dynamics and conformational changes at the binding pocket caused by the interaction with the ligand^17,33^. Further important advances have been made by grafting binding pockets between related PBPs^15,16^, although this has so far not expanded the spectrum to non-natural ligands. Therefore, we believe that our work is a crucial step forward and shows unmistakingly non-natural ligand-binding properties in RbsB mutants. Future studies on PBPs should focus on either improving experimental methods to select for better folders while maintaining designed or grafted new ligand-binding pockets, or on improving computational predictions of stability and translocation of designed mutants.

## Materials and Methods

### Computational design

For *in silico* design of the mutant library the Rosetta protein design software package was used. In particular, the ligand docking and the enzyme design modules within the Rosetta framework^18^ were used to predict amino acid changes in RbsB to potentially allow binding of 13CHD and CH. The protein design in Rosetta was carried out as a probabilistic simulated annealing algorithm for exploring the sequence space via rotamer replacement and optimization. The scheme has the following components: I-To parametrize and optimize the interaction via force-field terms; II-To determine the target residue positions to set the design and the one to repack; III-To iterate the cycles of sequence design and minimization; IV-To optimize structures using fixed rotamers without constraints. This defined the key residues in the RbsB binding pocket to be targeted (Table 1, Fig. 2) and computationally to be replaced by alanine. Subsequently, the 13CHD and CH molecules were docked 1000 times independently into the ‘“stripped” (Ala-substituted) RbsB binding pocket. The docked conformations were selected according to their energy values, and the ones with the lowest predicted energy values were used as a starting point for remodeling amino acid substitutions at the nine targeted positions. The final list of mutant positions was filtered according to the lowest minimal energy values, packing, hydrogen bond counts and ligand-solvent exposure.

### Mutant library and plasmid construction

Based on the *in silico* computational predictions, all the possible combinations of four alternate residues plus wild-type at nine positions (5^9^ combinations, Table 1) were produced by DNA synthesis as a mixture of linear DNA fragments (DNA2.0, USA). Delivered fragments were amplified by PCR with primers carrying tails that incorporated SalI and NdeI restriction sites. After restriction enzyme digestion and purification, the fragments were ligated with plasmid pSTV28P_AA_ ^12^ digested with SalI and NdeI, which brings expression of the *rbsB-* or its mutant gene under control of the P_AA_ promoter^34^ (Fig. S2). Multiple ligation reactions were independently transformed into batches of *E. coli* DH5α or MegaX-DH10B™ T1R Electrocomp™ cells (ThermoFisher Scientific). Transformants were cultured *en masse* on Luria-Bertani (LB) medium with chloramphenicol (Cm), from which the pool of pSTV28P_AA_-library plasmids was isolated and purified. Batches of 200 ng library-plasmid DNA were subsequently transformed into competent cells of the reporter strain *E. coli* BW25113 *ΔrbsB* containing plasmid pSYK1 (containing *trz1* under control of the LacI-repressed *tac-*promoter, and the *ompCp-gfpmut2* reporter; strain 4172)^12^ (Fig. S2, Table 2). Small proportions of these transformed batches were plated to estimate the number of viable clones in the libraries. The remaining pooled library cultures were grown for 16 h in 10 mL of low phosphate minimal medium (MM LP) (Table S1) containing 20 mM fumarate as sole carbon and energy source, and supplemented with Ampicillin (Ap) at 100 μg ml^−1^ and Cm at 30 μg ml^−1^ to select for both plasmids. Batches of 1.5 mL were aliquoted and stored in 15% (v/v) glycerol at −80°C.

Individual mutant clones selected from FABS screening (see below) were grown on LB plus Cm and Ap, and both plasmids (the pSTV28P_AA_-*rbsB-*mutant and pSYK1) were purified using NucleoSpin Plasmid columns (Machery-Nagel, Germany). Mutant *rbsB* genes were recovered on a fragment obtained by digestion with SalI and BstXI or XcmI, which was ligated into vector pET3d cut with the same enzymes^35^. This places the *rbsB* (mutant) gene with the hexahistidine tag at the C-terminal end under control of the T_7_ promotor, and removes the *rbsB* signal sequence for protein translocation to the periplasmic space. Ligations were transformed into *E. coli* BL21 (DE3) containing pLysS, for RbsB overexpression and purification (see below).

Two further RbsB mutant derivatives were produced individually by site-directed mutagenesis (DT002_H190N_ and DT016_M235V_) and one by site-saturation mutagenesis (DT016_W164G_), as follows. pSTV28P_AA_-derivative plasmids containing the respective mutant *rbsB* gene were amplified by PCR using overlapping but reverse complementary primers with point mutations at the desired positions. PCR products were digested with DpnI to remove template DNA^36^. After enzyme inactivation the PCR products were transformed into *E. coli* DH5α cells. Transformant colonies were selected on LB with Cm, plasmids were purified and the mutations in *rbsB* were verified by sequencing, after which they were transformed into the *E. coli* signaling reporter strain 4172 (see above)^12^.

### RbsB-based bioreporter assays

The capacity of RbsB or its mutants to induce the Trz1-OmpR *ompCp-gfpmut2* signaling chain in *E. coli* strain 4172 (Fig. S2) was assessed by flow cytometry, either on individual clones, uninduced or induced with the appropriate ligand in 96-well plates, or on mutant libraries with encapsulated cells grown to microcolonies in alginate microbeads, incubated as mixtures with or without inducer.

*E. coli* library aliquots of 50 μl (containing approximately 10^8^ cells) were inoculated in 10 ml MM LP medium (Table S1) containing 20 mM fumarate, and supplemented with 100 μg ml^−1^ Ap and 30 μg ml^−1^ Cm. Library batches were grown overnight at 37°C and with 180 rpm shaking. The next morning, cultures were diluted with MM LP to a turbidity of 0.03 and mixed in a 10:1 *v/v* ratio with 1% (*w/v*) alginate (PRONOVA UP LVG, FMC, Norway) in MM LP solution, to encapsulate the cells at approximately one starting cell per bead. Alginate beads were then formed using a VAR J30 bead machine (Nisco, Switzerland) at nozzle size of 150 μm and pressure set to 4-5 bar, and sprayed into 100 mM CaCl_2_ solution under constant stirring to solidify the alginate. This produces beads with an average diameter of 50 μm. After 1 h hardening in solution, cell-loaden beads were filtered sequentially through 40 and 70 μm mesh size nylon strainers (Corning Inc.) and washed with MM LP (Table S1). Recovered beads in the 40-70 μm-diameter range were then incubated for 16 h at 37°C in 5 ml MM LP containing 1 mM fumarate, Ap and Cm, in a rotating wheel (TC-7, New Brunswick/Eppendorf, Belgium). The next day, the cells had grown to microcolonies and individual beads were screened for fluorescence in a FACS Aria flow cytometer particle sorter (BD FACSAria Cell Sorter, Becton Dickinson, USA), equipped with a 100-μm nozzle at a flow rate of 2-5 μl s^−1^ and a density of between 100-1000 particles μl^−1^. Sensitivities for the FSC and FITC channels were set to 291 V and 435 V, respectively.

Microcolony-in-bead suspensions were screened first without induction and beads with fluorescence less than 1000 units were sorted to deplete the library of constitutive ‘ON’-mutants (Fig. 1A). Sorted beads were collected in LB medium supplemented with Ap and Cm, and incubated overnight at 37°C and with 180 rpm rotary shaking to dissolve the alginate beads and grow the *E. coli* cells. Multiple library batches were sorted sequentially to cover the entire mutant library.

The library depleted of constitutives was grown in multiple batches on MM with fumarate, cells were encapsulated and grown to microcolonies as described above, followed by 2.5 h induction with 1 mM 13CHD. Microcolony-in-bead fluorescence was again screened by flow cytometry, and beads displaying fluorescence levels higher than 1000 units were sorted in pools by FABS into tubes containing LB plus Ap and Cm medium (Fig. 1B). The collected beads were dissolved, regrown, and stored in 15% glycerol (*v/*v) at −80°C.

The resulting sub-libraries containing candidate 13CHD-responsive mutants were streaked on LB plates with Ap and Cm, and grown colonies were replica plated on MM agar with 20 mM fumarate, Ap and Cm (MM-FUM-ApCm) in presence or absence of 1 mM 13CHD. After 48 h incubation at 37°C the colonies were photographed under blue-light (Safe Imager Transilluminator, ThermoFisher) and their fluorescence intensities were compared by image analysis using the open source software: http://www.cheminfo.org/Image/Biology/Counting_plates/index.html?viewURL=https://couch.cheminfo.org/cheminfo-public/b616aba5eda653bf97ce9b776976aa4d/view.json?rev=37-e0a7762615ad2a544a1e5149ed1a2f21#). Colonies showing at least 1.25-fold increase in fluorescence in presence compared to absence of 13CHD were restreaked on LB-Ap-Cm agar plates and purified. Individual colonies were then inoculated in eightfold replicates in 96-well plates, containing per well 200 μl of MM-FUM-ApCm. 96-well plates were incubated overnight at 37°C and 700 rpm in a THERMOstar shaker (BMG LABTECH, Germany). The next morning, 5 μl culture of each well was transferred into a new well in a 96-well plate with 195 μl of fresh MM-FUM-ApCm. After 2 h incubation at 37°C, 100 μl from each well was transferred to the corresponding position of a new 96-well plate and immediately analyzed by flow cytometry to measure the uninduced fluorescence levels. To the remainder, 95 μl of fresh MM-FUM-ApCm, and 5 μl of inducer solution were added. This plate was incubated for another 2 h at 37°C, after which each well was again sampled for cellular fluorescence. As inducers we tested 0.1 mM ribose, 1 mM 13CHD and 1 mM CH (final concentrations in the assay). Cellular fluorescence was measured in 20 μl-aliquots, autosampled from each well by a Novocyte flow cytometer (ACEA Biosciences, USA), at an aspiration rate of 14 μl min^−1^ and culture density between 1000-5000 cells s^−1^. GFP fluorescence was recorded in the FITC-channel, which was set at a sensitivity of 441 V, and is reported as the average of the mean in each of the 8 replicates ± calculated standard deviations. Note that the fluorescence units of the FACS Aria (in Fig. 1) are not the same as the ones from the Novocyte (as in Table 4). Statistical significance was tested in pair-wise t-tests (one-sided, assuming increased response of the mutant).

### Expression and periplasmic space abundance analysis of RbsB wild-type or mutant proteins

The abundance of RbsB wild-type and mutants in the *E. coli* periplasmic space was analyzed using direct peptide mass identification, as described previously^12^. Periplasmic fraction was prepared by EDTA-ice treatment^12^ from *E. coli* BW25113 *ΔrbsB* carrying pSYK1 and the pSTVP_AA_-*rbsB* derivatives (Table 2). Periplasmic protein fractions were separated by SDS-PAGE and proteins in the size range between 28 and 36 kDa were excised from the gel. Proteins were analyzed by the UNIL Proteome Facility (https://www.unil.ch/paf/en/home.html). In short: samples were digested with trypsin and peptides were separated on an Ultimate 3000 Nano LC System (Dionex), followed by detection in a Thermo Scientific LTQ-Orbitrap XL mass spectrometer (Thermo Fisher Scientific, Waltham, MA). Mass spectra were analyzed using Scaffold Viewer 4, using thresholds of 99.9%. The minimum number of peptides for identification was 2.

### RbsB-His_6_ overexpression and purification

For purification of wild-type or mutant RbsB His_6_-tagged protein, 250 ml *E. coli* BL21 (pLysS) cultures with the corresponding pET3d-derivative plasmids grown in LB-Ap-Cm medium at 37°C until a culture turbidity at 600 nm of 0.3, were induced by addition of 1 mM isopropyl β-D-1-thiogalactopyranoside (IPTG, final concentration). Cultures were incubated further for 16 h at 20°C, after which the cells were harvested by centrifugation at 3,600 × *g* for 5 min at 4°C. Cell pellets were stored at −80°C until purification.

Thawed cell pellets were resuspended in 15 ml of buffer A (500 mM NaCl, 50 mM NaH_2_PO_4_, pH 8.0), containing 20 mM imidazole. The cell suspension was transferred to a metallic chamber containing a single metallic bead (50 ml chamber, Retsch, Germany) and frozen in liquid nitrogen for 1 minute. The cold chamber was transferred to a bead-beater machine (Oscillating Mill MM400, Retsch, Germany), and cells were crushed by constant vigorous shaking for 3 min at 30 s^−1^. The extract was transferred to a 50 ml centrifuge tube, which was centrifuged at 16,000 × *g* at 4°C for 30 min, after which the lysate supernatant was transferred to a clean tube.

The clean lysate was next loaded onto a HisTrap HP column (HisTrap FF crude 1 ml, GE Healthcare) at 4°C and flow rate of 0.5 ml min^−1^, followed by washes of, consecutively, 10 column volumes (cv, equal to 1 ml) of buffer A with 20 mM imidazole, 1.5 cv of buffer A with 40 mM imidazole and 1.5 cv of buffer A with 80 mM imidazole. Proteins were eluted with buffer A containing 250 mM imidazole in a total volume of 4 ml. The HisTrap eluate was subsequently loaded on a Superdex 200 10/300 GL 24 ml gel filtration column (GE Healthcare), and eluted with buffer A plus 250 mM imidazole at a flow rate of 0.75 ml min^−1^. Protein eluates were collected in aliquots of 150 μl, which were immediately frozen in liquid nitrogen and stored at −80°C, or used immediately for ITC assays. In case of RbsB mutant proteins we tested the effect of adding 10 mM ATP to the eluted protein solution directly after the HisTrap column, in order to disassociate and remove contaminating *E. coli* chaperones before loading onto the Superdex 200 10/30 GL column.

Protein concentrations were determined by NanoDrop spectrophotometry (Thermo Scientific, USA), using absorbance at 280 nm. The theoretical molar extinction coefficient and molecular weight were used as parameters. Subsamples of 20 μl were analyzed by SDS-PAGE to examine protein purity.

### Analysis of ligand binding using isothermal microcalorimetry (ITC)

A volume of 280 μl of purified protein extract (mostly the Superdex gel filtration eluate; between 5 and 10 mg protein ml^−1^) was pipetted into the measurement cell of a MicroCal ITC200 isothermal titration calorimetry instrument (GE Healthcare Life Sciences, USA). To avoid potential further unfolding of mutant protein we directly analysed them in imidazole-containing buffer A without previous dialysis, but maintained exactly the same volume of buffer A with 250 mM imidazole in the reference cell. An appropriate concentration of the test ligand (ribose or 13CHD; either at 0.1 or at 1 mM in buffer A with 250 mM imidazole) was filled into the injection syringe. Heat release was measured at 25°C with a reference power of 11 μcal s^−1^ and a stirring velocity of 1000 rpm. Raw data were recorded as changes in μcal s^−1^, and regression curves were fitted based on a one-binding site model using the Microcal Origin software (GE Healthcare).

### Circular dichroism and variable temperature measurements

Purified wild-type RbsB-His_6_, and DT002-His_6_ and DT016-His_6_-mutant proteins were analyzed by circular dichroism spectroscopy and variable temperature measurements using a J810 spectropolarimeter (Jasco, Japan). A volume of 100 μl of purified protein immediately after Superdex gel filtration or from thawed protein fraction stored at −80°C, was loaded on a PD minitrap G-25 column (GE Healthcare) and eluted with 500 μl of buffer A to remove the imidazole at 4°C. Protein in buffer A was then kept on ice and analyzed within 2 h for its circular dichroism spectrum.

Circular dichroism and thermal melting curves were determined in a quartz cuvette with a 0.1 cm path length (*L*). Spectra (θ, mdeg) were measured at room temperature between 200 and 260 nm at a scanning speed of 10 nm min^−1^ and a protein concentration of 0.1 mg ml^−1^. Buffer A alone was used as negative control and its circular dichroism spectrum was subtracted from that of the protein fractions. Data were further normalized for Δε (M^−1^ cm^−1^) using the effective protein concentration (*c*, mg ml^−1^) and the mean residue weight of RbsB (MRW, 109 Da), as follows:

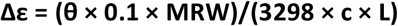

Circular dichroism spectra were further analyzed on the BeStSel webserver for secondary structure fold composition, as per instructions in Ref^24^.

Variable temperature measurements were conducted with a protein concentration of 0.3 mg ml^−1^ in buffer A (without imidazole) and the following parameters: Start temperature 20°C; temperature increment of 1°C; target temperature 95°C; temperature ramp rate of 2°C min^−1^. Measurements were performed in buffer alone or in presence of ribose at 0.1 mM or 13CHD at 1mM final concentration.

## Acknowledgements

The authors thank Luc Patigny and Julien Dénéréaz for their help in image analysis. The Gruber lab at the Department of Fundamental Microbiology is thanked for their help and advice in protein purification. This work was supported by grants 244405 (Biomonar) and OCEAN-2013-614010 (BRAAVOO) from the European Seventh Framework Program (FP7). Patrice Waridel and Manfredo Quadroni from the University of Lausanne Protein Analysis Facility are thanked for their help in peptide analysis.

## Author contributions

D.T., A.R., A. J. and V.S. performed experiments. S.R. and A.R. conducted computational designs. D. T., A. R. and J.R.M. prepared Figures. D.T., A.R. and V.S. contributed strains. D.T., A.R. and J.R.M. wrote the main manuscript. All authors reviewed the final manuscript.

## Additional Information

**Supplementary information** accompanies this paper at xxx

## Competing Interests

The authors declare that they have no competing interests.

## References

1. Chu, B. C. H. & Vogel, H. J. A structural and functional analysis of type III periplasmic and substrate binding proteins: their role in bacterial siderophore and heme transport. Biol Chem 392, 39 (2011).

2. Berntsson, R. P. A., Smits, S. H. J., Schmitt, L., Slotboom, D.-J. & Poolman, B. A structural classification of substrate-binding proteins. FEBS Lett 584, 2606–2617 (2010).

3. Quiocho, F. A. & Ledvina, P. S. Atomic structure and specificity of bacterial periplasmic receptors for active transport and chemotaxis: variation of common themes. Mol Microbiol 20, 17–25 (1996).

4. Li, H. Y., Cao, Z. X., Zhao, L. L. & Wang, J. H. Analysis of conformational motions and residue fluctuations for Escherichia coli Ribose-binding protein revealed with elastic network models. Intern J Mol Sci 14, 10552–10569 (2013).

5. Björkman, A. J. & Mowbray, S. L. Multiple open forms of ribose-binding protein trace the path of its conformational change. J Mol Biol 279, 651–664 (1998).

6. Dwyer, M. A. & Hellinga, H. W. Periplasmic binding proteins: a versatile superfamily for protein engineering. Curr Opin Struct Biol 14, 495–504 (2004).

7. Medintz, I. L. & Deschamps, J. R. Maltose-binding protein: a versatile platform for prototyping biosensing. Curr Opin Biotechnol 17, 17–27 (2006).

8. van der Meer, J. R. & Belkin, S. Where microbiology meets microengineering: design and applications of reporter bacteria. Nat Rev Microbiol 8, 511–522 (2010).

9. Looger, L. L., Dwyer, M. A., Smith, J. J. & Hellinga, H. W. Computational design of receptor and sensor proteins with novel functions. Nature 423, 185–190 (2003).

10. Baumgartner, J. W. et al. Transmembrane signalling by a hybrid protein: communication from the domain of chemoreceptor Trg that recognizes sugar-binding proteins to the kinase/phosphatase domain of osmosensor EnvZ. J Bacteriol 176, 1157–1163 (1994).

11. Srividhya, K. V. & Krishnaswamy, S. A simulation model of Escherichia coli osmoregulatory switch using E-CELL system. BMC Microbiol 4, 44 (2004).

12. Reimer, A., Yagur-Kroll, S., Belkin, S., Roy, S. & van der Meer, J. R. Escherchia coli ribose binding protein based bioreporters revisited. Sci Rep 4, 5626 (2014).

13. Reimer, A. et al. Complete alanine scanning of the Escherichia coli RbsB ribose binding protein reveals residues important for chemoreceptor signaling and periplasmic abundance. Sci Rep 7, 8245 (2017).

14. Schreier, B., Stumpp, C., Wiesner, S. & Höcker, B. Computational design of ligand binding is not a solved problem. Proc Natl Acad Sci USA 106, 18491–18496 (2009).

15. Banda-Vazquez, J. et al. Redesign of LAOBP to bind novel l-amino acid ligands. Protein Sci 27, 957–968 (2018).

16. Scheib, U., Shanmugaratnam, S., Farias-Rico, J. A. & Hocker, B. Change in protein-ligand specificity through binding pocket grafting. J Struct Biol 185, 186–192 (2014).

17. Boas, F. E. & Harbury, P. B. Design of protein-ligand binding based on the molecular-mechanics energy model. J Mol Biol 380, 415–424 (2008).

18. Leaver-Fay, A. et al. in Methods in Enzymology Vol. 487 (eds Michael L. Johnson & Ludwig Brand) Ch. 19, 545–574 (Academic Press, 2011).

19. Fleishman, S. J. et al. Computational design of proteins targeting the conserved stem region of influenza hemagglutinin. Science (New York, NY) 332, 816–821 (2011).

20. Chevalier, A. et al. Massively parallel de novo protein design for targeted therapeutics. Nature 550, 74–79 (2017).

21. Zoete, V., Cuendet, M. A., Grosdidier, A. & Michielin, O. SwissParam: A fast force field generation tool for small organic molecules. J Comp Chem 32, 2359–2368 (2011).

22. Bertelsen, E. B., Chang, L., Gestwicki, J. E. & Zuiderweg, E. R. P. Solution conformation of wild-type E. coli Hsp70 (DnaK) chaperone complexed with ADP and substrate. Proc Natl Acad Sci USA 106, 8471–8476 (2009).

23. Tokuriki, N. & Tawfik, D. S. Stability effects of mutations and protein evolvability. Curr Opin Struct Biol 19, 596–604 (2009).

24. Micsonai, A. et al. BeStSel: a web server for accurate protein secondary structure prediction and fold recognition from the circular dichroism spectra. Nucl Acids Res 46, W315–W322 (2018).

25. Vercillo, N. C., Herald, K. J., Fox, J. M., Der, B. S. & Dattelbaum, J. D. Analysis of ligand binding to a ribose biosensor using site-directed mutagenesis and fluorescence spectroscopy. Protein Sci 16, 362–368 (2007).

26. Yang, W. & Lai, L. Computational design of ligand-binding proteins. Curr Opin Struct Biol 45, 67–73 (2017).

27. Taylor, N. D. et al. Engineering an allosteric transcription factor to respond to new ligands. Nat Methods 13, 177–183 (2016).

28. Ray, S., Gunzburg, M. J., Wilce, M., Panjikar, S. & Anand, R. Structural basis of selective aromatic pollutant sensing by the effector binding domain of MopR, an NtrC family transcriptional regulator. ACS Chem Biol 11, 2357–2365 (2016).

29. Ko, W., Kim, S. & Lee, H. S. Engineering a periplasmic binding protein for amino acid sensors with improved binding properties. Org Biomol Chem 15, 8761–8769 (2017).

30. Reimer, A. Development of synthetic signaling pathways based on periplasmic binding proteins and hybrid chemoreceptors PhD thesis, University of Lausanne, (2017).

31. Duarte, J. M., Barbier, I. & Schaerli, Y. Bacterial microcolonies in gel beads for high-throughput screening of libraries in synthetic biology. ACS Synth Biol (2017).

32. Feldmeier, K. & Höcker, B. Computational protein design of ligand binding and catalysis. Curr Opin Chem Biol 17, 929–933 (2013).

33. Stank, A., Kokh, D. B., Fuller, J. C. & Wade, R. C. Protein binding pocket dynamics. Acc Chem Res 49, 809–815 (2016).

34. Alper, H., Fischer, C., Nevoigt, E. & Stephanopoulos, G. Tuning genetic control through promoter engineering. Proc Natl Acad Sci U S A 102, 12678–12683 (2005).

35. Studier, F. W., Rosenberg, A. H., Dunn, J. J. & Dubendorff, J. W. in Methods in Enzymology Vol. 185 (ed D. V. Goeddel) 60–89 (Academic Press, Inc., 1992).

36. Vogne, C., Beggah, S. & van der Meer, J. R. in Handbook of Hydrocarbon and Lipid Microbiology (ed K. N. Timmis) 4429–4444 (Springer, 2010).

