## Supplementary information for "Computational redesign of the *Escherichia coli* ribose-binding protein ligand binding pocket for 1,3-cyclohexanediol and cyclohexanol"

**Table S1-** Composition of mineral medium (MM) and low phosphate mineral medium (MM LP) used in this study.

| **Component** | **Mineral medium** | **Low phosphate mineral medium** |
|---|---|---|
| $Na_2HPO_4$ | 60 g | 0.36 g |
| $KH_2PO_4$ | 30 g | 0.33g |
| NaCl | 5 g | |
| $NH_4Cl$ | 10 g | |

Recipe for 1L. pH set to 7.4
Supplemented with 1 ml $l^{-1}$ of Hutner's trace mineral solution as per reference (1) for 21C medium.

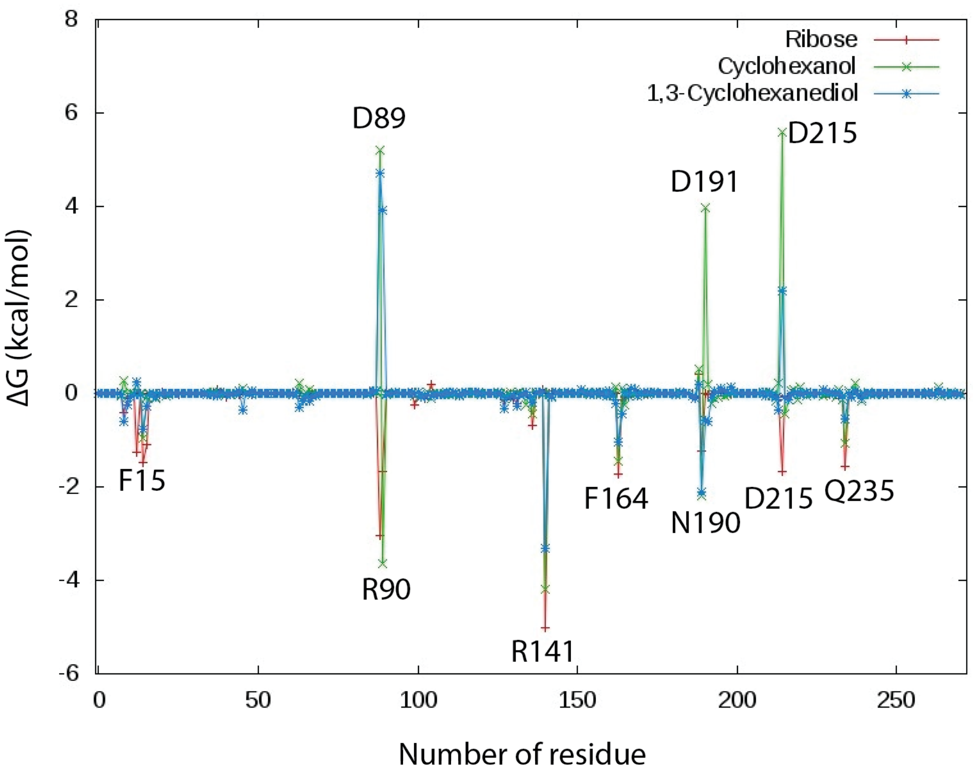


**Figure S1-** Contribution of each residue of RbsB to the change of Gibbs free energy ΔG (kcal/mol) during binding of the indicated ligand molecules using per-residue binding free energy decomposition based on Molecular Mechanics-Generalized Born Surface Area (MM-GBSA) method.

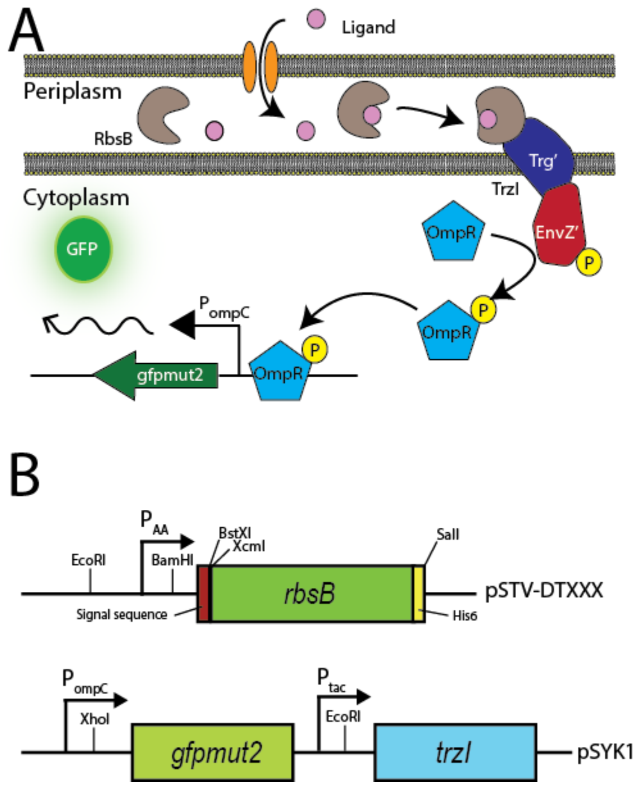


**Figure S2 -** The hybrid RbsB-OmpR signaling chain of the *E. coli* Trz1-OmpR bioreporter strain. (A) Ribose (ligand) is bound by the ribose-binding protein (RbsB), which docks to the Trz1 hybrid receptor (fusion between the periplasmic part of Trg and the cytoplasmic part of EnvZ). The binding starts a phosphorylation cascade, where OmpR is phosphorylated and increases transcription of gfp from the ompC promotor. (B) Scheme of relevant plasmids used in this work. Plasmid pSTV-DTXXX expresses the *rbsB* or mutant *rbsB* gene with its translocation signal sequence and hexahistidine tag ($His_6$) under control of the weak constitutive $P_{AA}$ promoter (2). Plasmid pSYK1 contains the *gfpmut2* gene under the *ompC* promoter control and the *trzI* gene under control of $P_{tac}$. Relevant restriction sites are indicated.

Supplementary References


1. **Gerhardt P, Murray RGE, Costilow RN, Nester EW, Wood WA, Krieg NR, Phillips GB (ed).** 1981. Manual of methods for general bacteriology. American Society for Microbiology, Washington, D.C.
2. **Alper H, Fischer C, Nevoigt E, Stephanopoulos G.** 2005. Tuning genetic control through promoter engineering. Proc Natl Acad Sci U S A **102:**12678-12683.
